# Glenn circulation causes early and progressive shunting in a surgical model of pulmonary arteriovenous malformations

**DOI:** 10.1101/2024.04.03.588015

**Authors:** Tina Wan, Henry Rousseau, Carol Mattern, Madeline Tabor, Matthew R. Hodges, Ramani Ramchandran, Andrew D. Spearman

**Author notes:** **Corresponding author:** Andrew D. Spearman, MD, 9000 West Wisconsin Avenue, Milwaukee, WI 53226.

## Abstract

**Background:** Pulmonary arteriovenous malformations (PAVMs) universally develop in patients with single ventricle congenital heart disease (CHD). Single ventricle PAVMs have been recognized for over 50 years, yet they are poorly understood, and we lack any medical therapies. To improve our understanding of single ventricle PAVM initiation and progression, we developed a surgical rat model of Glenn circulation and characterized PAVM physiology over multiple time points.

**Methods:** Using adult rats, we performed a left thoracotomy and end-to-end anastomosis of the left superior vena cava to the left pulmonary artery (unilateral Glenn), or sham surgical control. To assess for PAVM physiology in the left lung, we quantified intrapulmonary shunting using two independent methods (bubble echocardiography and fluorescent microsphere injection) at 2 weeks, 2 months, and 6 months. Additionally, we performed arterial blood gas measurements to assess oxygenation and plethysmography to assess ventilation.

**Results:** We identified pathologic intrapulmonary shunting by bubble echocardiography as early as 2 weeks post-Glenn surgery, and shunting continued chronically at 2- and 6-months post-Glenn. Shunting also progressed over time, demonstrated by increased shunting of 10µm microspheres at 6 months. Shunting was accompanied by mildly decreased arterial oxygenation, but there were no differences in ventilation as quantified by plethysmography.

**Conclusions:** Our surgical animal model of unilateral Glenn circulation re-creates the clinical condition of single ventricle PAVMs with early and progressive intrapulmonary shunting. This model is poised to characterize single ventricle PAVM pathophysiology and lead to mechanistic and therapeutic discovery.

**Graphic Abstract:** 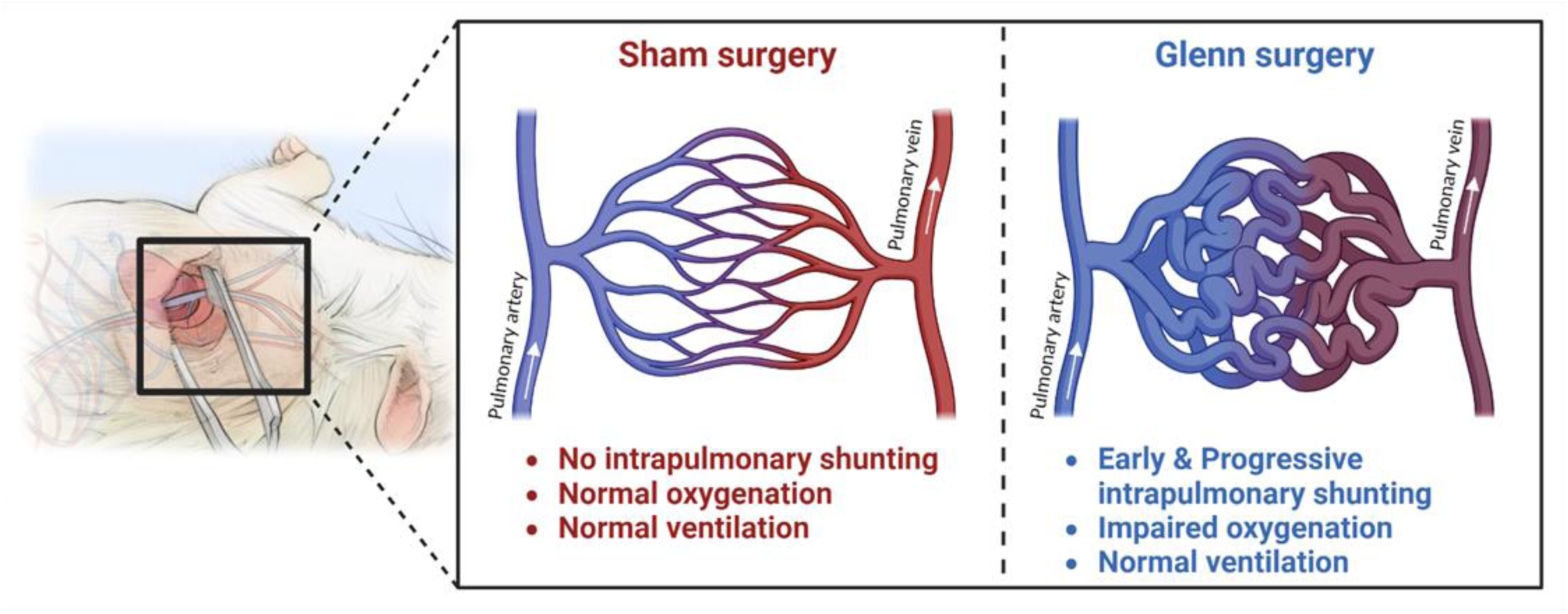

## Introduction

Pulmonary arteriovenous malformations (PAVMs) are vascular malformations that develop in hereditary and acquired conditions and lead to progressive hypoxemia [1, 2]. Among the acquired causes, single ventricle congenital heart disease (CHD) is perhaps the most well-recognized. Single ventricle PAVMs were first reported over 50 years ago when they were observed to develop after a surgical palliation known as the Glenn surgery [3, 4]. Since this initial report, there has been minimal progress in characterizing the fundamental features of single ventricle PAVMs, and there are no known medical treatments.

Clinical observations of single ventricle circulation support that PAVMs develop progressively after Glenn palliation, a surgical procedure where the superior vena cava (SVC) is directly anastomosed to a branch pulmonary artery and the main pulmonary artery no longer receives antegrade blood flow from the heart. As a result, pulmonary blood flow in Glenn circulation comes exclusively from the SVC, and venous blood from the lower body (including the inferior vena cava and hepatic vein) is pumped to the body via the single functional ventricle without perfusing the lungs [5]. In other words, PAVMs progressively develop when hepatopulmonary circulation is interrupted and pulmonary blood flow is delivered exclusively from the SVC.

Based on these strong observations of the importance of hepatopulmonary circulation and lack of *in vitro* PAVM models, previous research groups developed predominantly large animal models of unilateral right-sided Glenn circulation [6–9]. These studies demonstrated feasibility of establishing unilateral Glenn circulation in animals and validated that animal models of Glenn circulation develop pathologic intrapulmonary shunting as a sign of PAVMs. Additionally, one previous group even developed a small animal model of unilateral right-sided Glenn circulation in rats using microvascular surgery [10,11]. However, phenotyping of these animal models was limited, and the timing of PAVM initiation and progression in Glenn circulation remains poorly defined. Thus, we sought to modify this previously published rat model of Glenn circulation by performing a unilateral left-sided Glenn to take advantage of lung asymmetry and improve surgical visualization. As a first step to understanding single ventricle PAVM pathophysiology, we sought to characterize the timing of PAVM initiation and progression over multiple time-points in our rat model of Glenn circulation. To accomplish this, we used a combination of minimally invasive clinical diagnostic tools (bubble echocardiograms, arterial blood gases, and plethysmography) and an objective research technique (fluorescent microsphere injection). Collectively, we identified that our rat model of left-sided Glenn circulation re-creates the clinical condition of single ventricle PAVMs, which was demonstrated by early and progressive intrapulmonary shunting. Understanding the timing of PAVM progression is the necessary first step to mechanistic and therapeutic discovery.

## Methods

### Animals

Adult Sprague-Dawley rats (male and female, 6-12 weeks of age) were used for all experiments (Taconic). All rats were housed in the Biomedical Research Center at the Medical College of Wisconsin with access to standard chow diet, water ad libitum, and maintained in a 12-hour light/dark cycle. All surgical procedures described below were performed under isoflurane anesthesia (1-3%). All experimental protocols were approved by the Medical College of Wisconsin Institutional Animal Care and Use Committee prior to initiation of experimental protocols (Animal Use Agreement #7731).

### Left-sided superior cavopulmonary anastomosis (Left-sided Glenn)

To model Glenn circulation in rats, we performed an end-to-end unilateral cavopulmonary anastomosis between the left superior vena cava (L-SVC) and LPA. Our surgical approach is a modification of previous animal models of unilateral Glenn circulation, which were all previously performed on the right side (right SVC and right pulmonary artery (RPA)) [6, 9, 10]. We opted to perform a left-sided Glenn for two primary reasons: 1) improved visualization of the LPA compared to the RPA, and 2) the left lung consists of a single lobe versus the four lobed right lung, which allows continuity between different left lung regions as opposed to intrinsic regionalization of the right lung. Of note, unilateral Glenn circulation is possible in rats because normal venous anatomy in rats includes bilateral SVC. In contrast, normal venous anatomy in humans is a unilateral right SVC, and bilateral SVC is an uncommon variant in humans.

Prior to performing a left thoracotomy, rats were anesthetized with isoflurane, orally intubated, and mechanically ventilated (Rovent Jr, Kent Scientific) based on body weight. Buprenorphine ER (0.5 mg/kg, subcutaneous) was administered at the dorsal interscapular region, ophthalmic ointment was applied to the bilateral eyes to prevent corneal injury, and then rats were positioned in the right lateral decubitus position (left side up) on a warming pad. A pulse oximeter (Physiosuite, Kent Scientific) was attached to a lower extremity paw to assist physiologic monitoring during surgery. The surgical site was shaved with an electric shaver and the left chest was then cleansed with betadine followed by 70% ethanol (three times) and draped with clear sterile drape (Fig 1A).

**Figure 1.**
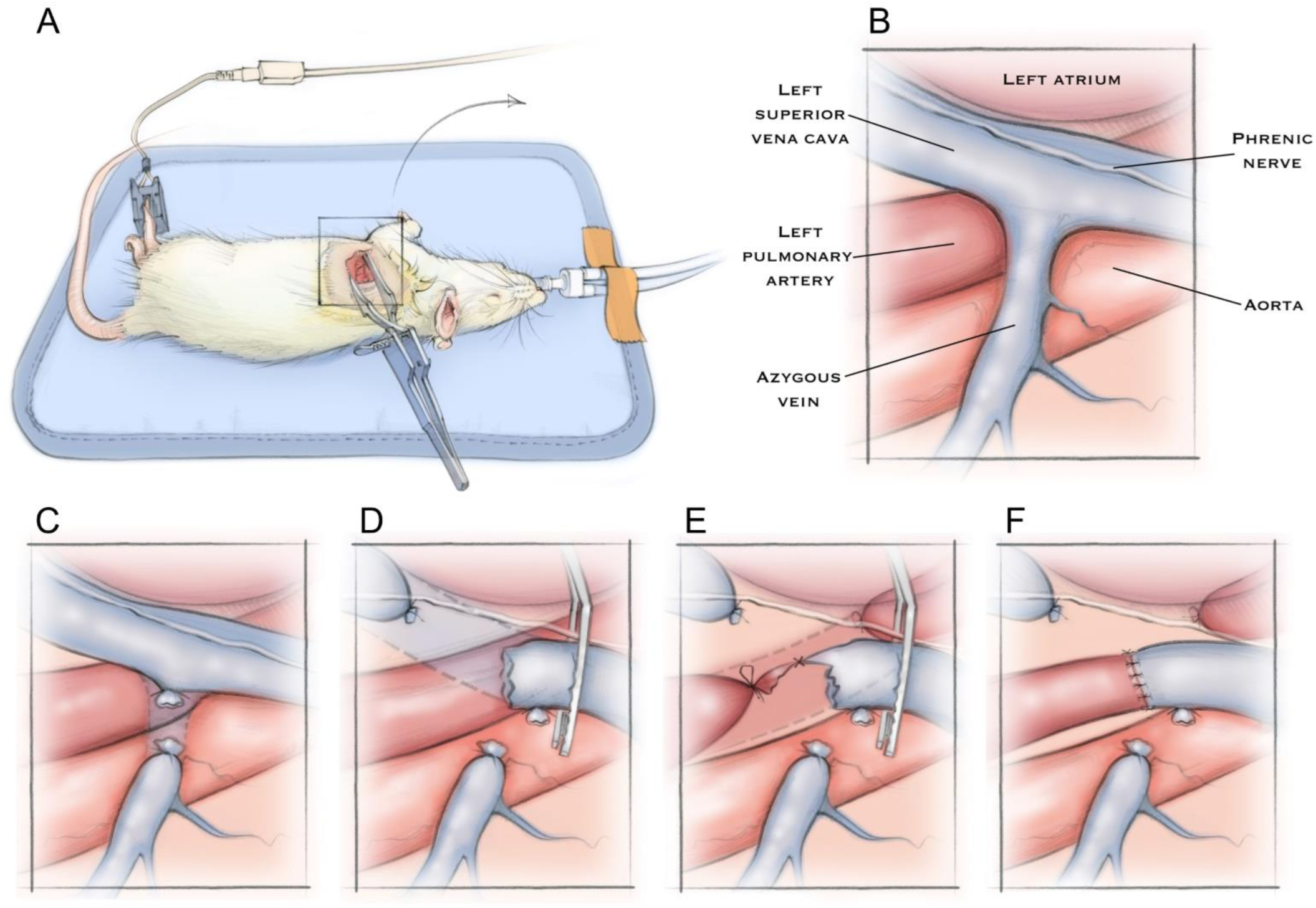
Rat positioning and surgical anatomy for left-sided end-to-end cavopulmonary anastomosis. (A) After intubation and initiating mechanical ventilation, rat is positioned in the right lateral decubitus position (left side up) for left thoracotomy, and a pulse oximeter is attached to a paw for physiologic monitoring during surgery. (B) Relevant structures visible in the surgical field after retraction of the left lung with sterile gauze. Step-by-step illustration of the cavopulmonary anastomosis includes (C) ligation and division of the azygous vein to prevent retrograde flow from the left superior vena cava (L-SVC) into the azygous vein. (D) Next, the L-SVC is dissected and isolated while preserving the overlying phrenic nerve. The distal L-SVC is tied off and a microvascular clamp is applied to the cranial aspect of the L-SVC. Then, the L-SVC is divided immediately cranial to the distal L-SVC suture, which allows exposure of the underlying left pulmonary artery (LPA). (E) The LPA is then dissected and isolated, the proximal LPA is suture ligated, a temporary slip knot is applied to distal LPA, and the LPA is divided. The end-to-end anastomosis between the L-SVC and LPA is constructed with a running suture posteriorly and interrupted sutures anteriorly. (F) After completing the anastomosis, the LPA slip knot and L-SVC clamp are removed to allow flow through the anastomosis. The chest is closed after appropriate hemostasis.

Left thoracotomy was performed at the 4^th^ intercostal space with the incision beginning ∼ 3 mm lateral to the sternum and extending ∼1.5cm. The chest wall was bluntly dissected, and the pleura opened to expose the left lung. The chest wall was then opened using an Alm self-retractor and the left lung was gently retracted from the surgical field using sterile gauze (Fig 1B). Using fine-tip jewelers forceps, the azygous vein was first dissected and isolated at its confluence with the L-SVC. The azygous vein was then ligated with 8-0 silk suture and divided (Fig 1C). The L-SVC was then dissected and isolated along its length to the coronary sinus while preserving the overlying phrenic nerve. An 8-0 silk suture was then tied at the distal aspect of the L-SVC to ligate the L-SVC from the coronary sinus. A microvascular clamp was then applied to the cranial aspect of the L-SVC, and the L-SVC was divided immediately cranial to the distal L- SVC suture (Fig 1D). After dividing the distal L-SVC, the LPA was widely exposed allowing dissection and isolation of the LPA. The proximal LPA was ligated immediately distal to the main pulmonary artery bifurcation with 8-0 silk. A temporary slip knot using 7-0 silk was applied to the distal LPA, and the LPA was divided immediately distal to the proximal suture. An end-to-end anastomosis between the L-SVC and LPA was then constructed using 9-0 nylon with a running suture along the posterior aspect and interrupted sutures anteriorly (Fig 1E). To aid visualization, sterile heparinized saline was applied to the surgical site and vessel lumens. After completing the anastomosis, the L-SVC clamp and LPA slip-knot were subsequently removed to allow flow through the anastomosis (Fig 1F). After appropriate hemostasis and visualization of flow through the anastomosis, chest wall was closed with a figure-of-8 suture (5-0 vicryl) followed by interrupted sutures (6-0 monocryl) to close the skin. Pre- and post-surgery videos of the surgical field are shown in Supplemental Videos 1 and 2.

After chest closure, isoflurane was stopped and ventilation was continued until the rat demonstrated consistent spontaneous ventilation, at which time the ventilation was stopped, and the endotracheal tube was removed. Animals recovered in a clean animal cage with overhead heating lamp until fully ambulatory.

For a comparative control animal, a similar sham procedure was performed. Sham rats were prepared in identical fashion to Glenn rats, including left thoracotomy and lung retraction with sterile gauze. The azygous vein was ligated and divided in identical fashion. The L-SVC was dissected and isolated in identical fashion, and a microvascular clamp was applied to the cranial aspect of the L-SVC for a similar duration as Glenn rats (∼20 minutes). The L-SVC clamp was then removed, and the chest wall and skin were closed in identical fashion. Glenn and sham rats were housed together in pairs of 2 or groups of 4 throughout the post-operative period.

### Bubble echocardiograms

To assess for intrapulmonary shunting on anesthetized rats, bubble echocardiograms were performed using transthoracic echocardiography (Vevo 3100, VisualSonics) and agitated saline injection into the L-SVC, similar to previous descriptions in humans, lambs, and pigs [8, 9, 12]. Rats were anesthetized with isoflurane and fur was removed from the chest with an electric shaver followed by depilatory cream. We then directly cannulated the left external jugular vein with a beveled catheter (MRE-027, Braintree Scientific), and the catheter was advanced ∼3cm into the L-SVC. Using a 3-way stopcock with two 5 ml syringes (3 ml saline in each syringe, 0.2 ml air in one syringe), we injected 0.5 ml agitated saline into the L-SVC with simultaneous visualization of the left and right ventricles by echocardiography. We semi-quantitatively assessed the severity of intrapulmonary shunting based on published clinical methodology [13–16]. Severity was quantified based on the subjective opacification (bubble density) in the left ventricle: negative (no identified bubbles), trivial (<10%), mild (10-50%), moderate (50-75%), and severe (>75%).

### Fluorescent microsphere shunting

Similar to a previous protocol using 6µm fluorescent microspheres to identify intrapulmonary shunting in a mouse model of PAVMs [17], we used 10µm fluorescent microspheres (ThermoFisher, F8834) to objectively quantify intrapulmonary shunting exclusively within the left lung. We used 10µm microspheres under the premise that these large microspheres cannot traverse the normally small caliber (< 5-10 µm) pulmonary capillaries, whereas PAVMs would permit pathologic shunting of microspheres through the lung parenchyma to the left atrium (i.e., intrapulmonary shunting).

Rat microsphere experiments were performed after intubation and ventilation, and immediately after euthanasia by exsanguination. The thoracic cavity was carefully opened to avoid injury to the lung. Ventilation was temporarily stopped to allow exposure to the distal LPA. The distal LPA was then carefully dissected at the hilum using the left mainstem bronchus as the inferior landmark. A beveled catheter (MRE-027, Braintree Scientific) was then inserted into the LPA ensuring that it was distal to the previous Glenn anastomosis in Glenn rats, or distal to the previous temporarily occlusion in sham rats. The catheter was advanced only 1-2 mm deep into the LPA to avoid wedging the catheter and assuring that all pulmonary artery branches were accessible with perfusion. The catheter was then secured with two 7-0 nylon sutures. After securing the catheter, ventilation was restarted to inflate the lungs. The left atrium was then widely opened to allow unrestricted pulmonary vein egress, and the left lung was gently perfused with 10 ml of PBS to remove blood and ensure full lung perfusion. We then injected ∼72,000 10µm fluorescent microspheres (20 µl microspheres diluted in 500 µl PBS) followed by 1 ml PBS flush. Microspheres and PBS flush were directly aspirated from the left atrium using a 1000 µl wide bore pipette tip (ThermoFisher, 2079GPK). Microspheres were then aliquoted into a 96 well plate and quantified using a fluorescent plate reader (SpectraMax i3x, ex/em 580/605). Shunt fraction was calculated as a percentage relative to the fluorescence of an equal number of non-injected microspheres: (collected spheres/non-injected spheres)*100.

### Arterial blood gases

To assess for potential differences in oxygenation, we collected arterial blood and measured the partial pressure of oxygen (PaO_2_), oxygen saturation (SaO_2_), and hemoglobin concentration (g/dL) using a point of care blood gas analyzer (iSTAT, Abbott; CG8+ cartridges, Medex Supply, ADE-03P8825). All animals were anesthetized with isoflurane, intubated, and mechanically ventilated according to body weight to ensure uniform ventilation. Additionally, because our unilateral Glenn model maintains a normal right lung, rats were ventilated with 100% oxygen to optimize the sensitivity for detecting potentially small differences in PaO_2_. All blood was collected free flowing from the right femoral artery using a 30g needle and heparin coated blood gas syringe (Fisher Scientific, 22-024-978).

### Ventilation

Non-invasive measurements of ventilation were made on conscious unrestrained rats in a custom-made 10-liter plexiglass whole body plethysmography, as previously described [18, 19]. Initially, rats were acclimated to the plethysmograph with a minimum of 3 training sessions of 20-30 minutes each (days 1-3). On day 4, baseline ventilatory function was then measured while breathing room air (FiO_2_=0.21) for 20 minutes, followed by a 10-minute hypercapnic challenge (FiO_2_=0.21, FiCO_2_=0.07). On day 5, baseline ventilatory function was re-measured while breathing room air for 20 minutes, followed by a 10-minute hypoxic challenge (FiO_2_=0.12). The following week, rats were re-acclimated for one day (day 8), and then hypercapnic and hypoxic challenges were repeated on subsequent days (days 9 and 10). Baseline, hypercapnia, and hypoxia measurements of breathing frequency (breaths per minute; bpm), tidal volume (VT; ml/breath/100g body weight), and minute ventilation (VE (ml/min/100g body weight) were averaged from week 1 and week 2.

### Statistical Analysis

Cohort data are expressed as mean and standard error of the mean (SEM) for continuous data and n (%) for categorical data unless otherwise stated. We used unpaired t-tests to evaluate the differences between Glenn rats and Sham rats at each time point, and one-way ANOVA to evaluate differences within groups at multiple time points. A p-value < 0.05 was considered statistically significant. Analyses were performed using GraphPad Prism 10 (GraphPad Software, San Diego, CA).

## Results

### Intrapulmonary shunting is detectable via bubble echocardiography early and chronically after surgery in Glenn rats

We first assessed for intrapulmonary shunting with a clinical diagnostic tool (bubble echocardiography) at 2 weeks, 2 months, and 6 months after surgery (Fig 2A-E). Bubble echocardiograms are performed under the premise that large caliber and transient air bubbles (agitated saline) are filtered out by the normal small caliber lung microvasculature and subsequently do not shunt through the lungs to reach the left ventricle. In contrast, large caliber air bubbles can shunt through the pulmonary vasculature in pathologic conditions such as PAVMs so that bubble are detectable by ultrasound in the left ventricle. In our rat model, we identified positive shunting in 5 of 7 Glenn rats (71% positive) as early as 2 weeks after surgery, whereas no sham surgery rats (0/6, 0% positive) had shunting reach the left ventricle (negative bubble echo) at 2 weeks. At 2 months, all Glenn rats had positive bubble echos (6/6, 100% positive), and all sham rats had negative bubble echos (0/6, 0% positive). Finally, at 6 months, we again observed that all Glenn rats had positive bubble echos (7/7, 100% positive). Interestingly, we did observe one sham rat at 6 months post-surgery had a trivial positive bubble echo (6 month sham: 1/4, 25% positive; total sham:1/16, 6% positive), which is in line with clinical studies that report ∼5% false positive bubble echos in healthy individuals [14]. Complete echocardiogram clips of sham and Glenn bubble echos are available in Supplemental Videos 3 (sham – negative), 4 (Glenn – moderately positive), and 5 (Glenn – trivially positive).

**Figure 2:**
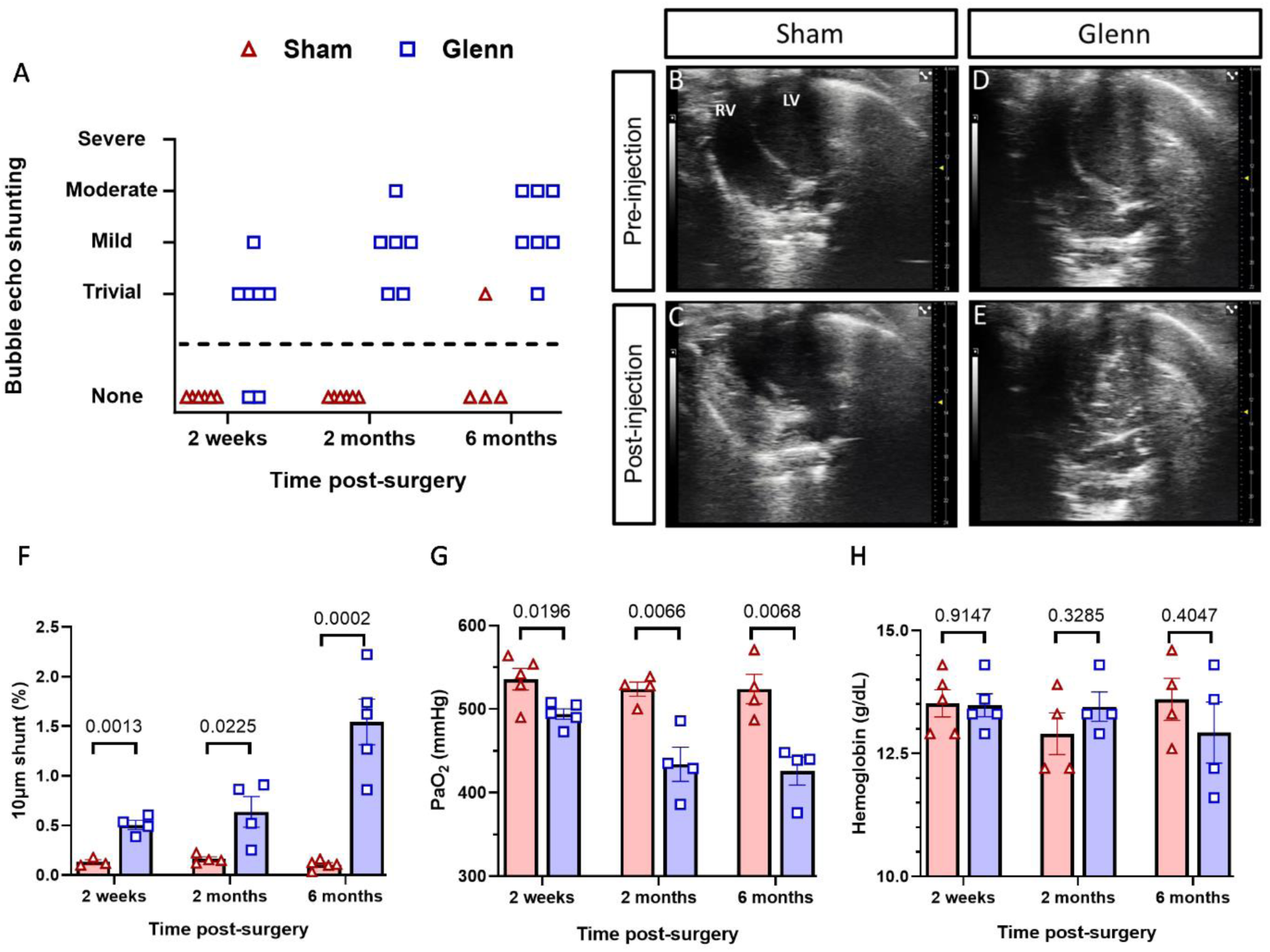
Glenn circulation leads to early and progressive intrapulmonary shunting with mildly decreased oxygenation. (A) Scatter plot of bubble echo shunting severity of sham control rats and Glenn rats at 2 weeks (n=6 sham, n=7 Glenn), 2 months (n=6 sham, n=6 Glenn), and 6 months (n=4 sham, n=7 Glenn) post-surgery. (B-E) Representative images of bubble echocardiograms. Images of sham and Glenn rat hearts pre-injection of agitated saline (B, D) show visualization of the right ventricle (RV) and left ventricle (LV). After injection of agitated saline (C, E), there is visualization of bubbles in the RV of sham rats but no bubbles in the LV (C), which indicates no intrapulmonary shunting. In comparison, there is visualization of bubbles in the LV of Glenn rats but no bubbles in the RV (E), which indicates pathologic intrapulmonary shunting. (F) Scatter plot with bars indicating mean and SEM for intrapulmonary shunting of 10µm microspheres. Shunt fraction (%) indicates the fluorescence of collected microspheres relative to fluorescence of non-injected microspheres. Unpaired t-tests at each time point (2 weeks: n=3 sham, n=4 Glenn; 2 months: n=4 sham, n=4 Glenn; 6 months: n=5 sham, n=5 Glenn). (G-H) Scatter plots with bars indicating mean and SEM for arterial partial pressure of oxygen (PaO_2_) and hemoglobin concentration measured by arterial blood gases performed with rats intubated, mechanically ventilated, and inspiring 100% oxygen. Unpaired t-tests at each time point (2 weeks: n=5 sham, n=5 Glenn; 2 months: n=4 sham, n=4 Glenn; 6 months: n=4 sham, n=4 Glenn).

### Intrapulmonary shunting of fluorescent microspheres progressively increases after surgery in Glenn rats

For an objective assessment of intrapulmonary shunting independent of bubble echocardiograms, we quantified intrapulmonary shunting of large caliber 10µm fluorescent microspheres by injection of microspheres into the distal left pulmonary artery (LPA) and direct collection from the left atrium. We identified increased shunting of microspheres in Glenn rats compared to their sham controls at every time point (2 week: p=0.001, 2 month: p=0.022, 6 month: p<0.001) (Fig 2F). Additionally, we observed that shunting progressively increased over time in the Glenn rats (2 week: 0.50±0.05%, 2 month: 0.64±0.15%, 6 month: 1.54±0.23%; one-way ANOVA p=0.003), whereas there was no difference in sham rats over time (2 week: 0.13±0.02%, 2 month: 0.16±0.02%, 6 month: 0.10±0.02%; one-way ANOVA p=0.22). These data are consistent with the clinical findings that PAVMs progressively increase over time in patients with Glenn circulation, which supports the clinical relevance of our model.

### Mildly decreased oxygenation in Glenn rats without systemic hypoxia

To assess for impaired oxygenation as a physiologic consequence of intrapulmonary shunting, we performed arterial blood gas measurements of PaO_2_ and SaO_2_. We identified decreased PaO_2_ in Glenn rats compared to sham controls at every time point (2 week: p=0.0196, 2 month: p=0.007, 6 month: p=0.007) (Fig 2G). Additionally, we again observed decreased PaO_2_ over time in Glenn rats (2 week: 494.2±6.2 mmHg, 2 month: 434.0±20.5 mmHg, 6 month: 425.8±16.7 mmHg; one-way ANOVA p=0.012), whereas there was no difference in sham rats over time (2 week: 535.8±12.9 mmHg, 2 month: 524.0±8.4 mmHg, 6 month: 524.0±17.7 mmHg; one-way ANOVA p=0.77). As expected, there were no differences in oxygen saturation with rats breathing 100% oxygen, and all SaO_2_ measurements were 100%.

Additionally, we measured hemoglobin concentration on all arterial blood gases to identify whether decreased PaO_2_ was associated with impaired oxygen delivery to tissues, which would be manifest with a compensatory increase in hemoglobin concentration. There were no differences in hemoglobin concentrations between Glenn and sham rats at any time point (2 week: p=0.91, 2 month: p=0.33, 6 month: p=0.40) (Fig 2H).

### No differences in ventilation measured via non-invasive plethysmography

Finally, we performed plethysmography to determine whether unilateral Glenn circulation impacts ventilation and breathing patterns under multiple conditions (room air [21% O_2_], hypercapnia [7% CO_2_], and hypoxia [12% O_2_]). As expected, there was no differences between Glenn and sham rats in breaths per minute, tidal volume, or minute ventilation at 2- and 6-months post-surgery under any condition (Supp Fig 1).

## Discussion

In this study, we demonstrate that surgical anastomosis of the L-SVC and LPA in rats (unilateral Glenn circulation) causes early and progressive intrapulmonary shunting indicative of PAVMs. Unilateral Glenn circulation re-creates clinical findings of progressive intrapulmonary shunting after Glenn palliation in patients with single ventricle CHD. Thus, this surgical rat model of Glenn circulation can be used to advance the field by characterizing the pathophysiology of single ventricle PAVMs. Additionally, this model has potential to identify critical components of hepatopulmonary interactions, including paracrine signaling from the liver that may regulate normal lung vascular development and homeostasis.

Previous groups reported developing surgical animals of unilateral Glenn circulation in pigs, lambs, and rats [6, 9, 10]. Similar to our study, they also reported universal development of intrapulmonary shunting in animals after undergoing unilateral Glenn surgery. Initial pig and lamb Glenn models performed bubble echos at 6-8 weeks post-surgery and reported positive versus negative shunting, as opposed to the semi-quantitative scale that is the clinical standard and the methodology we used here [6, 9]. More recently, McMullan et al performed bubble echos every week after surgery in lambs, and they reported consistent shunting (positive bubble echo) beginning at 5 weeks after surgery [8]. In contrast, the previously published right sided rat Glenn model assessed shunting with angiography, which is well-recognized to have low sensitivity for assessing PAVMs [20, 21]. Indeed, Starnes et al did not identify PAVMs in rats via angiography until 11 months after Glenn surgery [10]. Here, we rigorously assessed shunting over multiple time points in a small animal model of Glenn circulation, and we report the novel finding that shunting begins as early as 2 weeks after establishing Glenn circulation.

A critical clinical aspect of PAVM pathophysiology is that intrapulmonary shunting impairs oxygenation and leads to systemic hypoxemia. We identified that our unilateral model of Glenn circulation causes mild deficits in oxygenation, which was manifested by decreased PaO_2_ in arterial blood. Importantly, this mild decrease in PaO_2_ was accompanied by normal hemoglobin concentrations and no hemoglobin difference compared to sham counterparts, which suggests that Glenn rats did not have significant systemic hypoxemia. This is consistent with previous studies of unilateral Glenn circulation, including in pigs where Kavarana et al reported no differences in oxygen saturations [9]. Additionally, there were no differences in breathing under multiple conditions (baseline room air, hypercapnia, hypoxia), suggesting that Glenn circulation does not significantly alter baseline breathing patterns or normal chemoreceptor responses. Altogether, these findings support that unilateral Glenn circulation mildly impairs oxygenation; however, a healthy contralateral lung likely compensates to maintain normal systemic oxygenation. Thus, our model is not confounded by significant systemic complications.

Clinical observations support that the key factor driving PAVM formation in Glenn circulation is lack of hepatic vein perfusion to the lungs [2, 22–26]. It is unknown if other factors, such as non-pulsatile flow or biochemical mediators from SVC blood, are critical to the PAVM pathophysiology. Additionally, it is unknown how single ventricle PAVMs in Glenn circulation are similar or dissimilar to other clinical conditions causing PAVMs (hereditary hemorrhagic telangiectasia, Abernethy malformation) or intrapulmonary shunting (cirrhosis-induced hepatopulmonary syndrome). Given the phenotypic overlap of these conditions with single ventricle PAVMs, we speculate that our model of Glenn circulation will generate new insights into lung vascular development and homeostasis, physiologic paracrine signaling between the liver and lungs, and hereditary AVM pathogenesis. Thus, our model of Glenn circulation is highly applicable to diverse fields outside of pediatric cardiology and single ventricle CHD.

Despite the strengths of our model, this study has several limitations. First, lack of antegrade pulmonary blood flow to affected left lung, which is necessary for successful creation of this model, intrinsically leads to technical challenges when assessing intrapulmonary shunting. To overcome this, we reported two independent methods for assessing intrapulmonary shunting. Second, we assessed oxygenation in rats under isoflurane anesthesia, which may underestimate deficits in oxygen extraction compared to awake and ambulatory rats. Finally, six months post-surgery may not fully capture the long-term impact of unilateral Glenn circulation in rats. Later time points may be necessary to fully characterize vascular remodeling and potential systemic complications.

In conclusion, our surgical animal model of unilateral Glenn circulation re-creates the clinical condition of single ventricle PAVMs with early and progressive intrapulmonary shunting without secondary systemic complications. This model is poised to characterize single ventricle PAVM pathophysiology, as well as identify critical components of hepatopulmonary signaling that are likely essential for normal lung vascular development and homeostasis.

## Abbreviations

CHD: congenital heart disease
FiCO_2_: Fraction of inspired carbon dioxide gas
FiO_2_: Fraction of inspired oxygen gas
LPA: left pulmonary artery
L-SVC: left superior vena cava
PaO_2_: arterial partial pressure of oxygen
PAVMs: pulmonary arteriovenous malformations
RPA: right pulmonary artery
R-SVC: right superior vena cava
SaO_2_: arterial oxygen saturation

## Acknowledgements

Graphic abstract was created with BioRender.com.

## Sources of Funding

This study was supported by the National Institutes of Health from the National Heart, Lung, and Blood Institute (K08HL157510 - ADS), Medical College of Wisconsin Department of Pediatrics (ADS), and Herma Heart Institute Innovation Funds (ADS).

## Disclosures

None. The authors have no relationships with industry and no conflicts of interest.

## Author Contributions

Designing research studies-TW, MRH, RR, ADS

Conducting experiments-TW, HR, CM, MT, ADS

Acquiring data-TW, HR, CM, MT, ADS

Analyzing data-TW, RR, ADS

Writing and editing the manuscript-TW, HR, MT, RR, ADS

All authors approve the final version of this manuscript.

## Study Approval

All experimental protocols were approved by the Medical College of Wisconsin Institutional Animal Care and Use Committee prior to initiation of experimental protocols (Animal Use Agreement #7731).

## Supplemental Material

Supplemental Videos 1-5

Supplemental Figure 1

## Highlights

- Pulmonary arteriovenous malformations (PAVMs) are a universal vascular complication in single ventricle congenital heart disease that worsen pre-existing hypoxemia.
- This study reports development of a novel small animal model of Glenn circulation that phenocopies the clinical condition with development of early and progressive PAVMs.
- Mechanistic studies utilizing animal models of Glenn circulation are needed to define the pathophysiology of single ventricle PAVMs, identify therapeutic targets, and develop effective medical therapies.

## Notes

### Competing Interest Statement

The authors have declared no competing interest.

